# Habitat impacts the diversification of adhesive discs and skull shape in clingfishes

**DOI:** 10.64898/2026.09.07.749744

**Authors:** Jonathan M. Huie, Adam P. Summers, Kayla C. Hall, Thomas Trnski, Severine E. Hannam, Gary Voelker, Christopher M. Martinez, Kevin W. Conway

## Abstract

Specialized suction discs are functional innovations that enable fishes to attach to diverse surfaces and resist hydrodynamic forces. The adhesive discs of clingfishes (Gobiesocidae) vary in size and shape, but it is unclear how ecological factors have influenced their morphological evolution. Here we analyzed the disc and skull shape of 74 clingfish species using micro-CT scanning and 3D geometric morphometrics to investigate the role of habitat and substrate use on patterns of diversification. We also present novel comparisons of adhesive performance for 10 clingfish species. Clingfish interface directly with their environment using their adhesive discs, but we found that the disc and skulls are evolutionary integrated and share similar responses to habitat. Transitions from coastal habitats to coral reefs promoted elevated rates of evolution and morphological disparity across the body, whereas transitions to freshwater did not. Concurrently, repeated shifts from living on hard substrates to softer substrates (i.e., macroalgae and seagrass) were associated with more constrained disc shapes, convergent morphologies, and differences in adhesive performance. We propose that habitat and substrate use make complementary contributions to skeletal diversification in clingfishes, but the adhesive system requires further investigation to disentangle the complex interactions between form and function.

## INTRODUCTION

Habitat is a major axis of ecological variation that shapes patterns of morphological and functional evolution. Transitions to new habitats are met with shifts in selective pressures that may alter patterns of diversification (Betancur-R et al., 2012; Davis et al., 2012; Bouchenak-Khelladi and Linder, 2017). In aquatic environments, fishes are challenged with hydrodynamics forces acting to displace them. These forces vary across habitats and likely play an important role in the diversification of aquatic animals. High-flow habitats, including the rocky intertidal or fast flowing streams, appear to constrain evolution towards a narrow suite of phenotypes that reduce drag, such as streamlined body shapes and dorsoventrally flattened profiles in the case of benthic species (Langerhans, 2008; Lujan and Conway, 2015; Horn, 1999; Buser et al., 2017; de Barros and Caramaschi, 2019; Huie et al., 2019). In contrast, coral-reefs can provide structural relief from currents (Friedlander et al., 2003), and are often associated with faster rates of lineage and morphological diversification compared to non-reef habitats (Alfaro et al., 2007; Kiesling et al., 2010; Price et al., 2011). Because animals interact with their environment in diverse ways and have different strategies for coping with currents, habitat could have similar or variable effects on the evolution of morphological traits across the body.

Many aquatic animals resist displacement by currents by attaching to surfaces with diverse morphological structures, such as spines, friction pads, or suction organs (Ditsche and Summers, 2014; Delroisse et al., 2023). Specialized organs capable of generating suction have evolved independently across fishes in the form of ventral, dorsal, or oral adhesive discs (Friedman et al., 2013; Budney and Hall, 2010; Orr et al., 2019; Krings et al., 2023; Mcfarland et al., 2026). Adhesive discs likely evolved for station-holding in high-flow conditions, but they support a range of ecological niches. Gobies use their discs to scale waterfalls (Maie et al., 2012), remoras hitch a ride on megafauna (Flammang et al., 2020), and some snailfish adhere to crabs and deposit their eggs inside their host’s carapace (Yau et al., 2000). Given their ability to generate adhesion in diverse conditions, the adhesive discs of fishes have captivated biologists and engineers alike, serving as a source of inspiration for biomimetic suction cups (Gamel et al., 2019; Wang et al., 2017; Palecek et al., 2022; Wang et al., 2023). However, we still lack a general understanding of how habitat influences the morphology and evolution of adhesive discs at the macroevolutionary scale.

Clingfishes (family Gobiesocidae) have a well-developed ventral adhesive disc that provides an opportune system to investigate the relationships between ecology and morphology (Figure 1). There are nearly 200 clingfish species distributed around the world that occupy a range of marine habitats including the rocky intertidal, seagrass meadows, coral reefs, and deep subtidal waters to a depth of 560 m (Briggs, 1955; Hofrichter and Patzner, 2001; Fricke et al., 2017; Conway et al., 2020). A few species even occupy brackish and freshwater systems (Conway et al., 2017a,b). We hypothesize that habitats impose different demands on clingfishes, with high-flow environments requiring better adhesion and more flattened cranial profiles.

**Figure 1.**
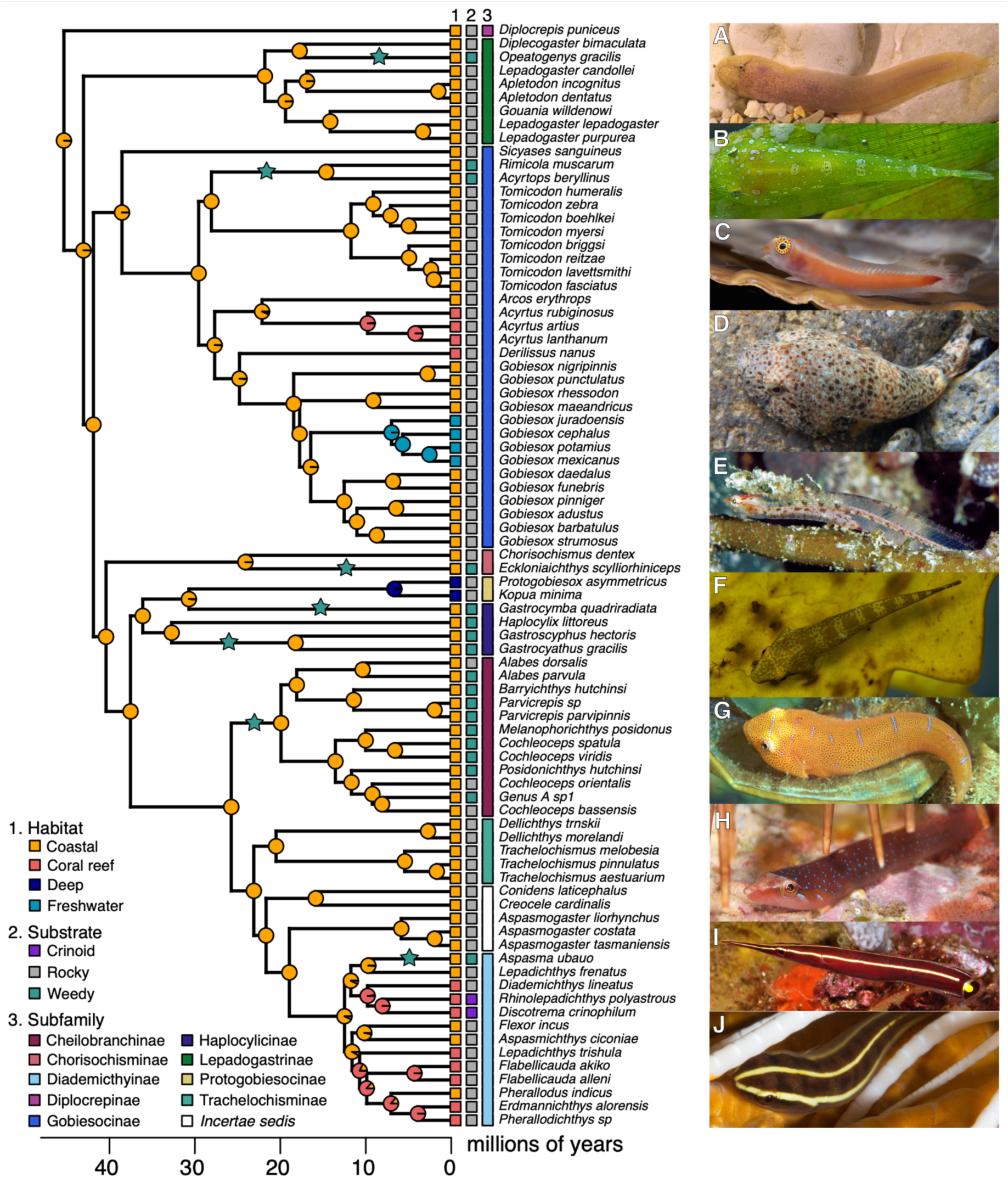
Time-calibrated phylogeny of 82 clingfish species. Phylogeny depicts the ancestral state reconstruction of habitat, with habitat, substrate use, and subfamily distributed across the tips. Green stars indicated branches where transitions from rocky to weedy substrates likely occurred (see Figure S2). Live photos highlight clingfish diversity. A) *Gouania sp.* (photo by M. Wagner), B) *Acyrtops beryllinus*, C) *Derilissus sp* (photo by B. Brown/coralreefphotos). D) *Gobiesox cephalus*, E) *Alabes parvula* (photo by R. Kuiter), F) *Parvicrepis parvipinnus*, G) *Cochleoceps orientalis* (photo by *G. Short*), H) *Dellichthys morelandi* (photo by I. Skipworth), I) *Diademichthys lineatus* (photo by M. Erdmann), and J) *Discotrema crinophilum* (photo by M. Erdmann).

Within their respective habitats, clingfishes are also found in association with different substrate types. While clingfishes typically dwell on hard “rocky” substrates (i.e., boulders, rubble, shells, hard coral), species spanning multiple subfamilies have evolved specialized “weedy” lifestyles on the surfaces of macroalgae or seagrass (Briggs, 1955; Roland 1978; Hofrichter and Patzner, 2001; Conway et al., 2019; Conway et al., 2024). Clingfish discs may have evolved to improve attachment to the substrates that they primarily live on. This is evidenced by the rocky specialist *Gobiesox maeandricus*, which adheres better to rough and hard surfaces than to smooth and flexible ones (Wainwright et al. 2013; Ditsche et al. 2014; Huie and Summers 2022). Weed-dwelling species also share physical characteristics including narrow, elongate bodies and relatively narrow heads that likely reflect adaptations for living on comparably smooth macroalgae and seagrass blades (Conway et al., 2019; 2024). This suggests that interactions between habitat and substrate use may influence the patterns of diversification across the clingfish adhesive disc and skull.

Clingfish adhesive discs are a complex structure, made up of elements from both the pectoral and pelvic girdles (Figure 2). On the contact surface, the disc features papillae, small hierarchical pads, that putatively improve suction by increasing hydrodynamic adhesion and friction (Wainwright et al., 2013; Ditsche and Summers, 2019; Sandoval et al., 2020; Hernandez et al., 2025). The performance of one clingfish species (*G. maeandricus*) is well-documented. It can generate adhesive forces up to 250 times its body weight, adhere to rough and fouled surfaces, and outperform lumpsucker and snailfish species that also have adhesive discs but occupy deeper habitats with weaker hydrodynamic demands (Green and Barber 1988; Wainwright et al. 2013; Ditsche et al. 2014; Huie et al., 2022). Biomimetic suction cups based on clingfish discs suggest the bones play an important role in supporting the disc and preventing internal collapse, while a soft disc rim deforms to create a tight seal on irregular surfaces (Ditsche and Summers, 2019; Sandoval et al., 2019). Across the family of clingfish, the adhesive discs vary greatly in size and can span the entire belly, cover a comparatively small proportion of the belly, or be completely absent as in *Alabes* (Springer and Fraser, 1976). The skeletal morphology of the disc is also highly variable (Briggs, 1955), yet it is unknown how this morphological diversity or its relationship with adhesive performance factors into an ecomorphological framework.

**Figure 2.**
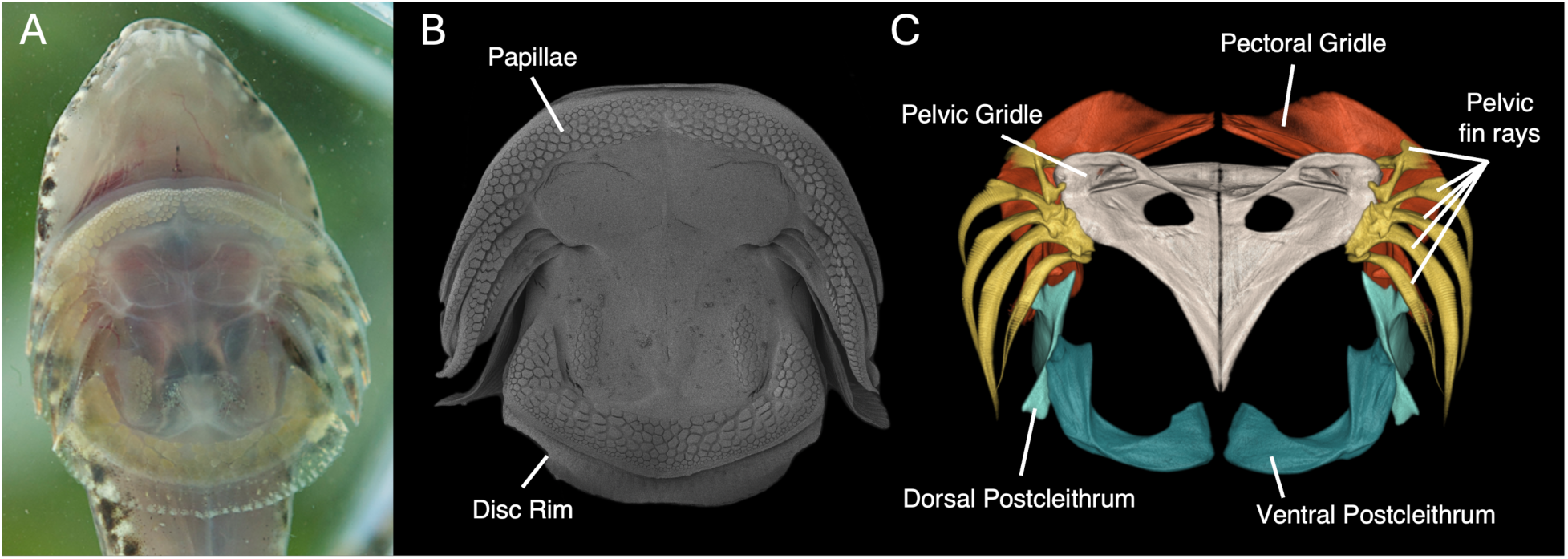
Images of the clingfish adhesive disc. A) Ventral photo of *Gobiesox cephalus*. showing its large adhesive disc. B) Scanning electron micrograph of the adhesive disc in *G. cephalus*. C) Rendering of a micro-CT scan of the disc skeleton.

Here, we examined the evolutionary relationships between ecology, morphology, and adhesive performance in clingfishes. We present a 3D geometric morphometric dataset using micro-CT scans for 74 clingfish species in 42 genera as well as novel performance data for 10 species. We examined the skeletal elements of the disc and the skull, and used phylogenetic comparative methods to analyze the tempo and mode of their evolution. Our aims were two-fold: 1) investigate the effects of habitat and substrate use on clingfish morphology, phenotypic disparity, and rates of evolution and 2) compare the adhesive performance of clingfishes with different substrate preferences. We predicted that transitions away from high-flow coastal environments, which are expected to impose strong constraints, would be associated with increases in phenotypic disparity and evolutionary rates in the disc and skull. Additionally, we predicted that substrate use may have a stronger effect on disc shape than skull shape.

Specifically, the repeated evolution of weed-specialists may have promoted convergence in disc shape and led to differences in adhesive performance between species living predominantly on weedy or rocky substrates.

## MATERIALS AND METHODS

### Phylogeny and divergence time estimates

We inferred a time-calibrated phylogenetic hypothesis for Gobiesocidae by reanalyzing data from Conway et al. (2020). The previous study proposed relationships between 82 extant clingfish taxa but not their divergence times. We used calibrations employed in a previous study of clingfishes (Conway et al., 2017a) based on outgroup relationships derived from Near et al. (2013). Currently, only two fossil clingfishes have been described, but they are not ascribable to any extant species and therefore cannot be attributed to an internal node (Schwarzhans et al., 2017; 2024).

The time calibrated tree was estimated using BEAST v1.10.4 (Drummond et al., 2012). We used seven partitions, according to the PartitionFinder results of Conway et al. (2020) and equal to the number of genes in their dataset. We unlinked both substitution model and clock model and linked all tree models. Following Rüber et al. (2020), we changed the following priors from their default value: clock rate for the seven genes were changed to Gamma (1, 1), initial = 1; and all GTR substitution parameters were changed from Gamma to InverseGamma. We enforced monophyly on seven groups (following the results of Conway et al., 2017a, 2020), including: (1) Blennidae; (2) *Entomacrodus nigricans* + *Salarias fasciatus*; (3) Blennidae + Gobiesocidae; (4) all gobiesocids excluding *Diplocrepis puniceus*; (5) subfamily Gobiesocinae (New World gobiesocids); (6) core group of *Gobiesox* including four freshwater (*G. cephalus, G. juradoensis, G. mexicanus,* and *G. potamius*) and six marine species (*G. barbatulus, G. strumosus, G. pinniger, G. adustus, G. daedaleus,* and *G. funebris*); and (7) the four freshwater species of *Gobiesox*. We applied lognormal calibration priors to four of the seven aforementioned monophyletic groups (1–3, 6) based on inferred divergence times from Conway et al. (2017). This included: (i) mean age of 36.2 million years (standard deviation 2.5, offset 1.0) for the most recent common ancestor (TMRCA) of Blennidae (group 1 above); (ii) mean age of 32.4 million years (standard deviation 2.5, offset 1.0) for TMRCA of *Entomacrodus nigricans* + *Salarias fasciatus* (group 2); (iii) a mean age of 69.0 MY (standard deviation 2.0, offset 1.0) for TMRCA of Blennidae + Gobiesocidae (group 3); and (iv) a mean age of 13.5 million years (standard deviation 2.8, offset 1.0) for TMRCA of core group of *Gobiesox* (group 6).

We then ran two separate uncorrelated relaxed molecular clock analyses in BEAST, under the Yule process. Markov chain Monte Carlo (MCMC) chains were run for 200 million generations, sampling every 20,000 generations. The log files resulting from each of the two analyses were evaluated in Tracer v1.7.2 (Rambaut et al. 2018), to measure effective sample size (ESS) and to determine the number of generations to discard as burn-in, and then combined using LogCombiner v1.10.4. The tree files resulting from each of the two analyses were also combined in LogCombiner v1.10.4 and summarized into a single maximum clade credibility (MCC) tree using TreeAnnotator v1.10.4 (Drummond et al., 2012).

### Ancestral state reconstruction of ecological traits

We reconstructed the evolutionary history of habitat and substrate use across clingfishes. The 82 species sampled in the phylogeny were assigned to one of four habitat categories encountered across the family: coastal, coral reef, freshwater, or deep-water (Table S1). The coastal environment encompassed intertidal and seagrass inhabitants, while coral reefs included species further offshore. The freshwater category included only obligate freshwater species and the deep-water species were those with a maximum depth greater than the epipelagic zone (200 m). No deep-water species were available for sampling in the morphological analyses. Species were also classified into one of three substrate categories: rocky, weedy, or crinoids (Table S1). Rocky substrates included hard structures like boulders, gravel, shell hash, or hard coral; while weedy substrates encompassed macroalgae and seagrass. The crinoid category consisted only of the two genera of specialists that dwell on crinoids (sea lilies). While species of the genus *Alabes* lack an adhesive disc, one species (*Alabes parvula*) still exhibits strong affinity for weedy substrates and was coded as such, while the other (*A. dorsalis*) was coded as a rock-dweller. One caveat with our approach is that clingfishes may use multiple habitats or adhere to diverse substrates throughout a lifetime and do not perfectly conform to these categories (e.g., the cleaner clingfish *Cochleoceps orientalis* establishes cleaning stations on both rocky and weedy substrates; Hutchins, 1991). In such cases, we opted for the most commonly-associated substrate of a given species.

We reconstructed the joint evolution of habitat and substrate preference with hidden Markov models using the *CorHMM* R package v2.8 (Boyko and Beaulieu, 2021; Beaulieu et al., 2022). We fit three transition rate models (equal rate, symmetric, and all-rates different) with and without a hidden state parameter, and all with and without simultaneous transitions between habitat and substrate for a total of 12 models. We generated 1,000 stochastic character mappings using parameters from the best fitting model based on weighted AIC scores.

### Morphological sampling and shape analyses

To quantify the shape of the skulls and bony elements of the adhesive disc, we obtained micro-computed tomography (μCT) scans for 74 extant clingfish species (1 specimen per species) representing 38% of described species and nine of the ten subfamilies. Sampling was conducted to maximize the coverage of species sampled across the phylogeny. A few species not represented in phylogeny (*Derilissus nanus* and *Lepadichthys frenatus*) were sampled as substitutes for congeners in the phylogeny but for which morphological specimens were not available (*D. lombardii* and *L. misakius*). Specimens were primarily scanned at the University of Washington Friday Harbor Laboratories, with one scan done at Harvard’s Museum of Comparative Zoology and another at the Natural History Museum, London. Additional scans were obtained from MorphoSource (Boyer et al., 2016; Blackburn et al., 2024). A comprehensive list of sampled specimens and their voucher information are presented in Table S2.

Skeletal morphology was quantified using 3D geometric morphometrics. We used 3D Slicer and the SlicerMorph extension to visualize and place landmarks on the volumetric renderings of the scans (Rolfe et al., 2021). We digitized 229 landmarks (107 fixed and 122 sliding semi-landmarks) across the left side of the disc and skull (Table S3; Figure S1). Within the disc, we analyzed three structures: the pelvic girdle (basipterygium), pectoral girdle, and ventral postcleithra. The pelvic girdle and ventral postcleithra provide structural support for the adhesive disc, while the pectoral girdle articulates with the pelvic girdle and provides attachment points for muscles. These structures are invariably present across clingfishes, except for *Alabes* that have secondarily lost their disc and their pelvic girdle and postcleithra are either absent or vestigial (Springer and Fraser 1976). To handle the missing disc structures in the two species of *Alabes* in our dataset, we landmarked the skulls and pectoral girdle as normal but the pelvic girdle and ventral postcleithral were each represented by repeating landmarks placed in a single location where the disc would have been (Bardua et al., 2019). For the skull, our landmarking scheme matched Neves et al. (2025) and spanned eight structures: the neurocranium, premaxilla, maxilla, dentary, angular-articular, hyomandibula, hyoid bar, and urohyal. These bones hold functional importance during the diversification of fishes with regards to feeding, respiration, and housing sensory organs (Larouche et al., 2023; Buser et al., 2024; Neves et al., 2025).

After digitization, we imported the landmark coordinates into R Statistical Environment version 4.4.1 (R Core Team, 2024). Because sampling only one side of bilaterally symmetric structures can skew shape variation along the midline (Cardini, 2016; Bardua et al., 2019), we mirrored the landmarks on neurocranium and pelvic girdle from the left side onto the right side across a midline plane using the “mirrorfill” in the paleoMorph R package (Lucas, 2016; https://cran.r-project.org/package=paleomorph). We then accounted for non-shape variation in size, translation, and rotation of the specimens by performing a Generalized Procrustes Analysis on the whole dataset (global superimposition) with the “gpagen” function in the *geomorph* R package (Baken et al., 2021). Semi-landmark points were allowed to slide while minimizing Procrustes distances. Due to fish skeletons being highly kinetic, we standardized the positions of the different bones using a local superimposition (Rhoda et al., 2021). Separate Generalized Procrustes Analyses were performed on each bone individually; then the locally aligned coordinates were translated to the average position and scaled to the average size of the corresponding bone in the global superimposition performed on the whole dataset. Following the local superimposition, we used principal component analyses to reduce data dimensionality and visualize the primary axes of skeletal variation. Because the absence of disc elements in *Alabes* greatly distorted the axes of the morphospace, we separated the locally superimposed dataset into a skull dataset with all 74 species and a disc dataset with 72 species without *Alabes.* These datasets were analyzed separately in all subsequent analyses.

### Testing for ecomorphological relationships and convergence

We investigated the effects of habitat and substrate use on the variance and average trends in skeletal shape. First, we performed phylogenetic MANOVAs using the “procD.pgls” function in *geomorph* to compare mean shapes between the habitat and substrate categories.

Size was included as a covariate in these analyses to account for potential effects of allometry. Because local superimpositions complicate the use of centroid sizes for an entire configuration, we use the log-transformed centroid sizes of pelvic girdle and neurocranium to represent disc and skull size, respectively (shape ∼ log(centroid size) * ecology). Additionally, we compared the amount of phenotypic disparity associated with each habitat and substrate category using the “morphol.disparity” function in *geomorph*. For MANOVAs and disparity analyses, significance was based on 1,000 iterations. To complement these analyses, we also estimated phylogenetic signal within the disc and skull datasets to assess the extent to which phylogeny explains the patterns of variation. We used the “phylosignal” function in *geomorph* to estimate the multivariate extension of Blomberg’s K (K_mult_) for each dataset with 1,000 simulations for significance testing.

Given that clingfishes are expected to have repeatedly evolved the use of weedy substrates, we investigated whether these taxa exhibit morphological convergence in disc and skull shapes using two methods. First, we estimated Stayton’s C metrics of convergence (Stayton, 2015) using the “convSig” function in the “convevol” R package (Brightly and Stayton, 2024). We calculated C_1_ which measures how close focal taxa are to each other in the morphospace relative to their ancestors. Convergence tests were performed on each dataset with 1,000 simulations. Additionally, we tested for convergence using the “search.conv” function in the *RRphylo* package (Castiglione et al., 2019). This method compares angles (θ_real_) between phenotypic vectors between focal taxa as well as θ_real_ divided by evolutionary time separating focal taxa; in both cases smaller angles represent greater similarity. For both sets of convergence analyses, we used PC scores as the input trait data. Because choosing how many PC axes to retain can be arbitrary, we used two different sets of PC scores. First, we performed the analyses using the first 14 PC axes of the disc morphospace and first 12 PC axes of the skull morphospaces, which represented 85% of the total shape variation in each respective dataset.

Since many of those axes explained proportionally small amounts of the total variation (< 2%), we used visual Cattell scree tests to retain 4 and 5 PC axes from the disc and skull datasets, which represented 66% and 68% of the total variation, respectively.

### Tempo of skeletal evolution

To help explain patterns of disparity, we compared the net rates of skeletal evolution between habitat and substrate groups. We used the multivariate Brownian Motion framework implemented by the “compare.evol.rates” function in *geomorph*. We also used the “compare.multi.evol.rates” function in *geomorph* to evaluate differences in the net rates of shape evolution between the disc and skull. *Alabes* specimens were omitted for this comparison. For both rate analyses, the inputs were the superimposed Procrustes coordinates and significance was tested based on 1,000 simulations.

We also tested for shifts in evolutionary rates across the clingfish phylogeny. We fit two models of evolution using BayesTraitsV4 (Pagel and Meade, 2022), a single rate Brownian motion model that assumes one rate across the phylogeny, and a variable rates model that allows for changes in rates throughout the phylogeny and identifies where rates differ (Venditti et al., 2011). Inputs of the analyses were the PC axes representing 85% of the total variation in each dataset. Following Evans et al. (2023), we multiplied the PC scores by 1,000 to correct for known computational difficulties arising from small numbers in BayesTraits. Evolutionary correlations between PC axes were accounted for using the “TestCorrel” function in BayesTraits. We then used a reversible-jump Markov chain Monte Carlo method with uniform priors and ran two independent chains for 200 million generations, sampling every 10,000 iterations, with the first 60 million discarded as burn-in. Convergence was assessed by rerunning the analysis and visually inspecting the trace of the marginal likelihoods using Tracer v1.7.2 (Drummond et al. 2012). Model comparison was done by calculating Bayes factors from the marginal likelihoods of the models, with values greater than 10 regarded as strong support for the variable rates model.

### Comparing adhesive performance

We compared the adhesive performance across 10 coastal species of clingfish, including nine species collected from the North Island of New Zealand for this study and *Gobiesox maeandricus* from the Pacific Northwest (Table S4). Collection of New Zealand fishes were allowed under Fisheries New Zealand Special Permit number 691 issued to Auckland Museum by the Ministry of Primary Industries. For each New Zealand specimen, we recorded the standard length, disc surface area, and body mass (n = 1-14 specimens per species). We measured the force required to pull freshly euthanized specimens off hard substrates of varying roughness following the procedures of Wainwright et al. (2013) and combined our data with their data for *G. maeandricus* (n = 22 specimens). Briefly, the specimens were attached to submerged substrates made from epoxy resin with roughnesses that corresponded to six different grit sizes (0, 15.3, 35, 52, 78 and 127 μm). Medical suture (5-0, 6-0, or 7-0 hypalon) was looped through the body of the fish and used to attach it to the cross-head of a material testing system (MTS). The MTS was used to pull the fish at a speed of 1 mm sec−1 until it detached from the substrate. Prior to each test, we pressed down on the fish to evacuate water under the disc and ensure adhesion. Specimens were tested on some or all of the substrates between one and 10 times.

In this study, we only analyzed data for one surface roughness. We used 15.3 μm because the most number of specimens were tested on this substrate (n = 58). For each specimen, we retained the maximum force recorded across trials. Because adhesive forces are dependent on disc size, which varies with overall body size and between species, we normalized peak force by dividing by disc area; resulting in peak adhesive stress.

We used linear mixed effect models (LMMs) to assess variation in adhesive performance across clingfishes using the *nlme* R package v3.1 (Pinheiro et al., 2025). First, we compared the peak stress between rock-dwelling (n = 8) and weed-dwelling (n = 3) species, using substrate as the fixed effect and individual as the random effect [Stress ∼ Substrate + (1|ID)]. To explore the effects of substrate use on disc effectiveness, we compared the relationships between adhesive force and disc area [log(Force) ∼ log(Disc Area) * Substrate + (1|ID)]. Then we investigated variation in peak stress across species [Stress ∼ Species + (1|ID)]. To compare point estimates of peak stress, we calculated the estimated marginal means (EMMs) and standard error for each of the fixed effects (species or substrate) using the emmeans R package v1.10 (https://CRAN.R-project.org/package=emmeans).

### Evolutionary integration of shape and performance

We investigated the degree of evolutionary integration between morphological structures and performance in several ways using the “phylo.integration” function in *geomorph*. First, we investigated the covariance between disc and skull shape across all clingfishes. *Alabes* were omitted from these analyses and significance testing was performed using 999 iterations. Second, we also estimated the evolutionary covariance between adhesive performance and disc shape.

Estimated marginal means of peak stress and whole disc shape for ten species were used as the inputs. Third, we also estimated the integration between peak stress and each of the elements of the adhesive disc independently.

## RESULTS

### Evolutionary history of habitat and substrate use

The evolutionary changes in habitat and substrate use in clingfishes were best described by symmetric transition rates without simultaneous state changes between the two traits. The ancestor of modern clingfishes most likely inhabited a coastal environment (98.5%) and adhered to rocky substrates (94.9%) (Figure 1; S2). We recovered four transitions away from coastal environments to coral reefs, and one reversal. On coral reefs, there was one transition from rocky substrate to associations with crinoids in the ancestor of *Discotrema* and *Rhinolepadichthys* (Figure S2). There was also a single transition to deep waters in the ancestor of *Kopua* and *Protogobiesox*. Consistent with a previous study, we recovered a single transition to freshwater habitats within *Gobiesox* (Conway et al. 2017). Lastly, we recovered seven independent transitions from rocky substrates to weedy substrates and at least four reversals back, all within the coastal environment (Figure 1; S2).

### Axes of variation in skeletal morphology

Clingfishes displayed a wide range of disc shapes, with a large proportion of species clustered in one quadrant of the morphospace (Figure 3A, 3B). Here, the pelvic girdle was narrower with a modest dorsal crest and the ventral postcleithra were more paddle-like. There was an offshoot from the main cluster towards the left side of the first principal component (PC 1; 41.3% of the variation), primarily represented by the genus *Gobiesox*. These species had broad pectoral girdles with laterally expanded cleithra, flatter and wider triangular pelvic girdles, and large ventral postcleithra that were more “L” shaped. A second offshoot from the main cluster occurred at low PC 2 scores (11.1% of the variation) and was represented by members of the subfamily Diademichthyinae. These species had relatively larger pectoral girdles and smaller pelvic girdles. The pelvic girdles were shorter with taller dorsal crests and less triangular in shape. Instead, these girdles were more laterally expanded in the posterior region where they articulated with the ventral postcleithra.

**Figure 3.**
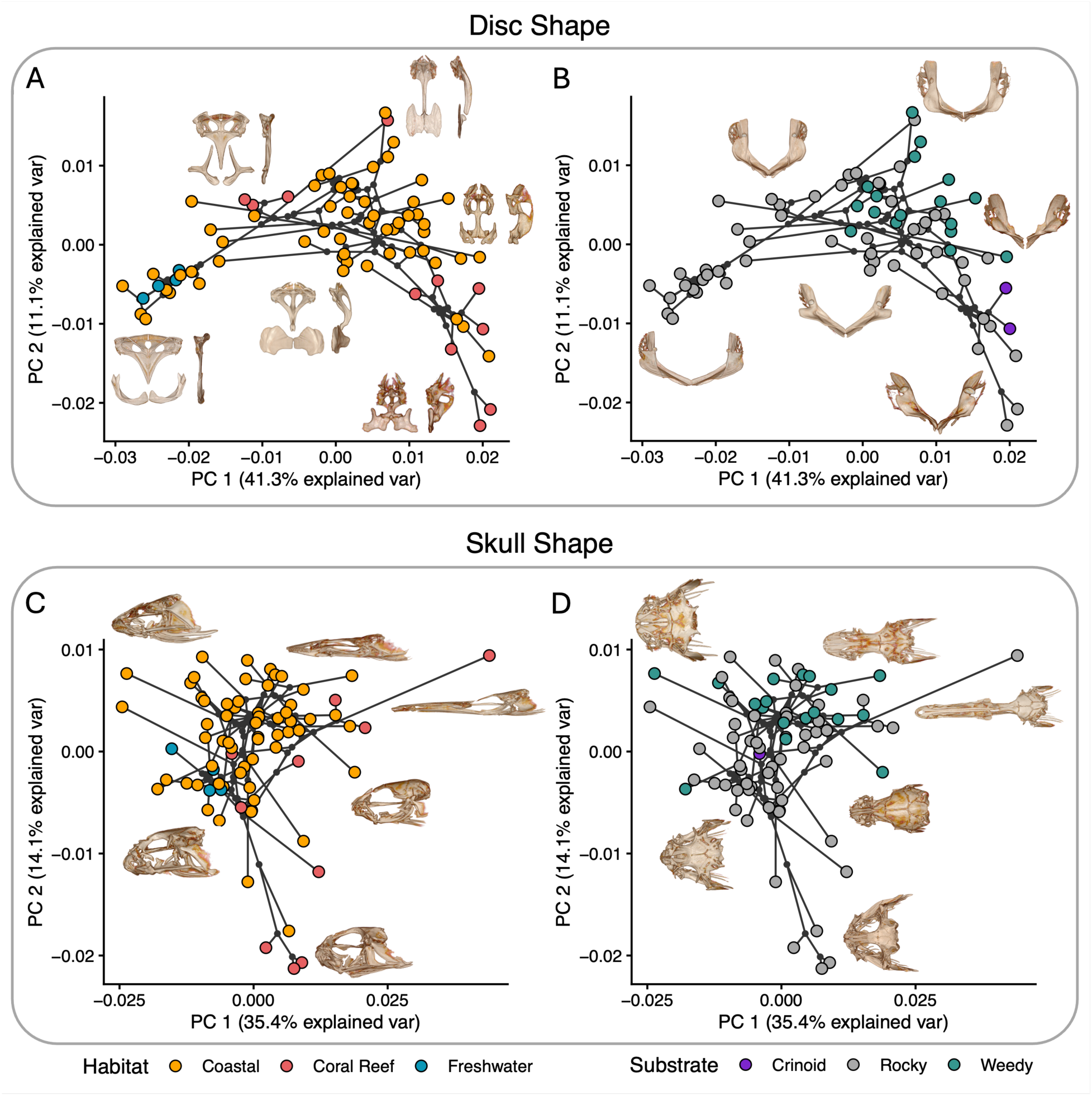
Phylomorphospaces depicting variation in clingfish disc and skull shapes. A) and B) show the same disc morphospace colored by habitat and substrate use, respectively. Insets depict exemplar variation in the pelvic girdle and ventral postcleithra from the ventral and lateral perspective (A) or an anterior view of the pectoral girdle (B). C) and D) show the same skull morphospace colored by habitat and substrate use, respectively. Insets depict exemplar skulls from the lateral (C) and dorsal (D) views.

Clingfish skulls also exhibited highly varied phenotypes, but were more evenly distributed than the discs, except for one morphologically extreme species, *Diademichthys lineatus* (Figure 3C, 3D). PC 1 of the skull morphospace (35.4% of the variation) largely reflected variation in the aspect ratio of the skull. Most species occupied the region of PC 1 associated with relatively stout skulls, while *D. lineatus* exhibited a long, narrow, and flattened skull. PC 2 (14.1% of the variation) largely reflected differences in head height, tapering of the snout, and orbit size. Most species had fairly dorsoventrally flattened skulls, broad snouts, and small to moderately sized orbits. In contrast, species of the genera *Acyrtus* and *Arcos* had taller skulls, pointed snouts, and substantially larger orbits. Across the family, clingfish disc and skull shapes exhibited significant evolutionary integration (r-PLS = 0.729, p = 0.003). Pelvic girdles with taller dorsal crests were generally associated with narrower skulls and smaller orbits.

Estimates of phylogenetic signal indicated that disc shape exhibited a stronger phylogenetic signal (K = 0.760, p = 0.001) than skull shape (K = 0.511, p = 0.001).

### Associations between ecology and morphology

Habitat and substrate contributed to variation in phenotypic disparity and evolutionary rates, but not differences in mean shape (Figure 4). Phylogenetic MANOVAs found no significant effect of size or ecology on shape (Table S5). Coastal species encompassed almost the full extent of the shape variation in both morphospaces, but exhibited similar levels of disparity as coral reef species (p ≥ 0.065). However, similar disparity levels were achieved at different paces depending on habitat; the discs and skulls of coral reef species evolved 2x faster than the coastal species (p ≤ 0.001). Freshwater species exhibited significantly less disparity than coastal and coral reef species and evolved the slowest.

**Figure 4.**
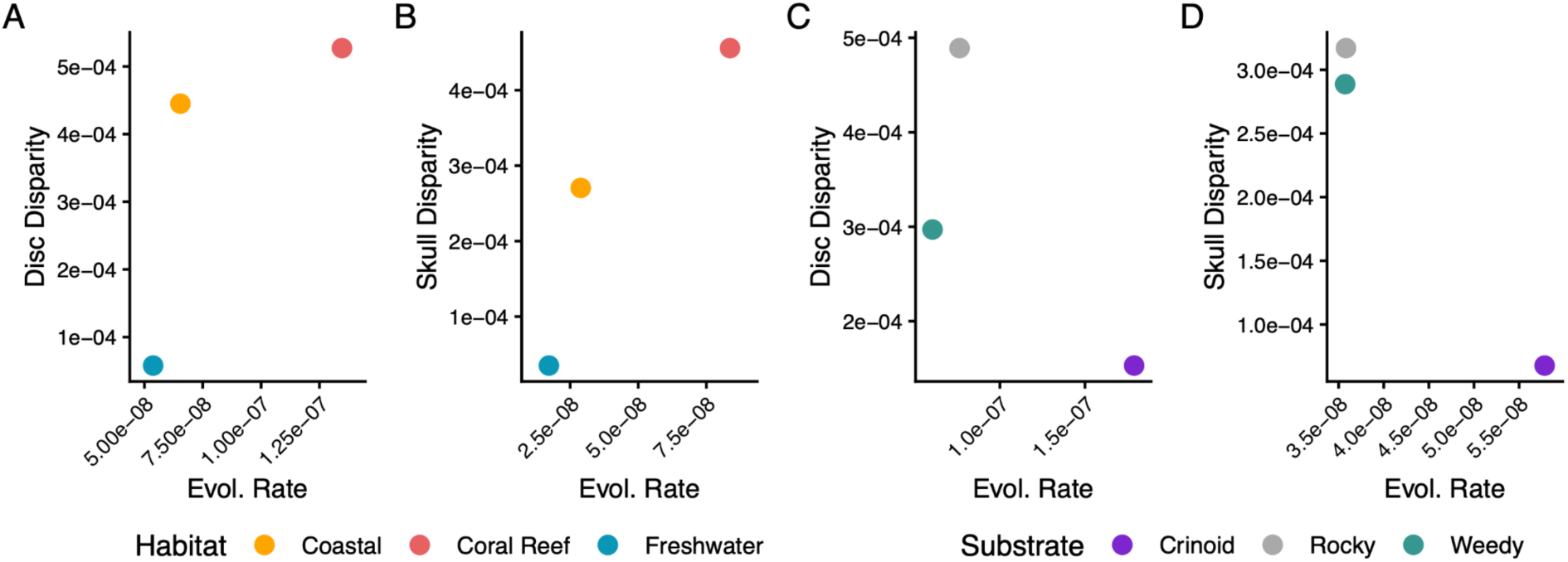
Phenotypic disparity and evolutionary rates of the disc and skull shape of gobiesocid fishes. Panels are colored by habitat (A, B) and substrate use (C, D).

Substrate use had no significant effect on the disparity (p ≥ 0.100) or evolutionary rates (p = 0.286) of skull shape, but impacted disc shape. Rock-dwelling species, often coinciding with coastal habitats, also encompassed the full range of shape variation and exhibited 1.6x greater disc shape disparity than weed-dwellers (p = 0.005). Differences in disc shape disparity were attributed to differences in evolutionary rates, as rock-dwellers evolved 1.2x faster than those of weed-dwellers (p = 0.001). Crinoid species appeared to exhibit the lowest levels of disparity but the fastest rates of evolution, though we advise caution with these results as they are based on just two species.

Analyses of phenotypic convergence support weed-dwellers as having converged on similar skeletal shapes (Table 1). Analysis of Stayton’s C found evidence for significant convergence in skull shape with both the more inclusive and truncated set of PC axes.

**Table 1.** Results of the weed-dweller convergence analyses.

| | C1 | C1 p-value | $\theta_{real}$ | $\theta_{real}/time$ | $\theta_{real}$ p-value | $\theta_{real}/time$ p-value |
| --- | --- | --- | --- | --- | --- | --- |
| Disc - 14 PCs | 0.056 | 0.342 | 69.810 | 1.156 | **0.001** | **0.003** |
| Disc - 4 PCs | 0.170 | **0.020** | 56.860 | 0.911 | **0.001** | **0.001** |
| Skull - 12 PC | 0.112 | **0.001** | 78.283 | 1.391 | **0.001** | 0.526 |
| Skull - 4 PC | 0.177 | < **0.001** | 76.543 | 1.363 | **0.001** | 0.464 |

Meanwhile, weed-dwellers exhibited significant convergence in disc shape with only the smaller PC trait matrix. This suggests that weed-dwellers exhibit convergence along the major axes of variation, but maintain differences along axes that explain smaller amounts of variation. Further support for convergence in the disc and skull was provided by the RRphylo convergence tests.

Both the skulls and discs of weed-dwellers exhibited significant convergence with both sets of PC scores. However, the convergence in the skull was not significant when accounting for the amount of evolutionary time separating weed-dwelling taxa.

### Tempo of shape evolution

Net rates of skull shape evolution were 2x faster than disc shape (p = 0.008), but both phenotypes were described by a variable rate Brownian motion model of evolution (Figure 5). In terms of disc shape, we found a large rate increase in the ancestor of *Gobiesox* as well as along the branch leading to the weedsucker, *Eckloniaichthys*. There were multiple rate shifts within Diademichthyinae, specifically in the ancestor of *Fabellicauda* and the common ancestor of *Diademichthys* and the two crinoid specialist genera. Diademichthyinae and the sister clade (*Aspasmogaster*) also exhibited high and variable rates of skull shape evolution, with *Diademichthys* exhibiting the fastest tip rate of any clingfish. The subfamily Cheilobranchinae, which includes many of the Australian weed-dwellers, also exhibited faster rates of skull shape evolution, particularly the disc-less *Alabes.* Elevated rate shifts were also observed in the ancestors of *Acyrtus* and *Arcos* as well as *Gouania wildenowi*, an interstitial specialist. Finally, branches leading to the two largest species (*Chorisochismus dentex* and *Sicyases sanguineus*) and to one of the smallest (*Derlissus nanus*) all exhibited elevated rates of skull evolution.

**Figure 5.**
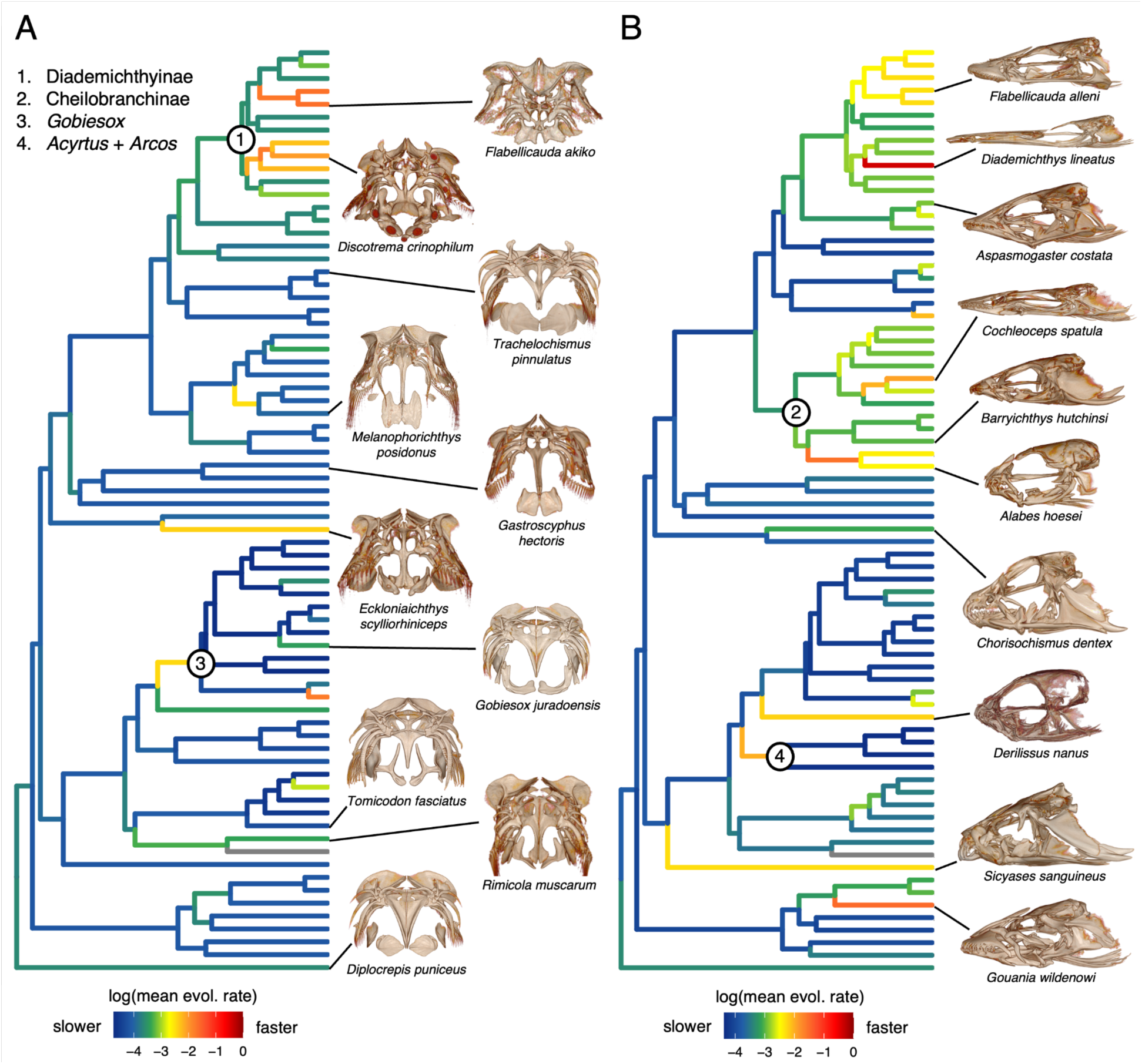
Tempo of skeletal shape evolution across the clingfish phylogeny. Estimated rate shifts for A) disc shape and B) skull shape evolution were mapped onto the phylogeny. Warmer colors represent faster rates and cooler colors represent slower rates. Insets show representative micro-CT images of the disc and skull.

### Clingfish adhesive performance

Peak adhesive stress varied significantly among species (F = 3.609, p = 0.006) and between rock and weed-dwellers (F = 13.930, p < 0.001) (Figure 6). On average, the adhesive discs of rock-dwellers generated 1.5x more adhesive stress than weed-dwellers on the hard substrate. This was partially explained by the relationships between adhesive force and disc area (Figure 6B). Rock-dwellers exhibited a higher intercept (F = 15.054, p = 0.002) and steeper slope (F = 9.207, p = 0.003) than weed-dwellers, resulting in rock-dwellers generating more adhesive force per unit of disc area than weed-dwellers. Rock-dwellers were sampled from distinct regions of the original disc morphospace, but point estimates of maximum adhesive stress varied minimally between 33.6 and 41.3 kPa, except for *Trachelochismus aestuarium* that averaged 25.8 kPa (Table S4; Figure 6C, 6D). In contrast, weed-dwellers ranged from 14.2 to 29.7 kPa and clustered in the same region of the morphospace (Table S4).

**Figure 6.**
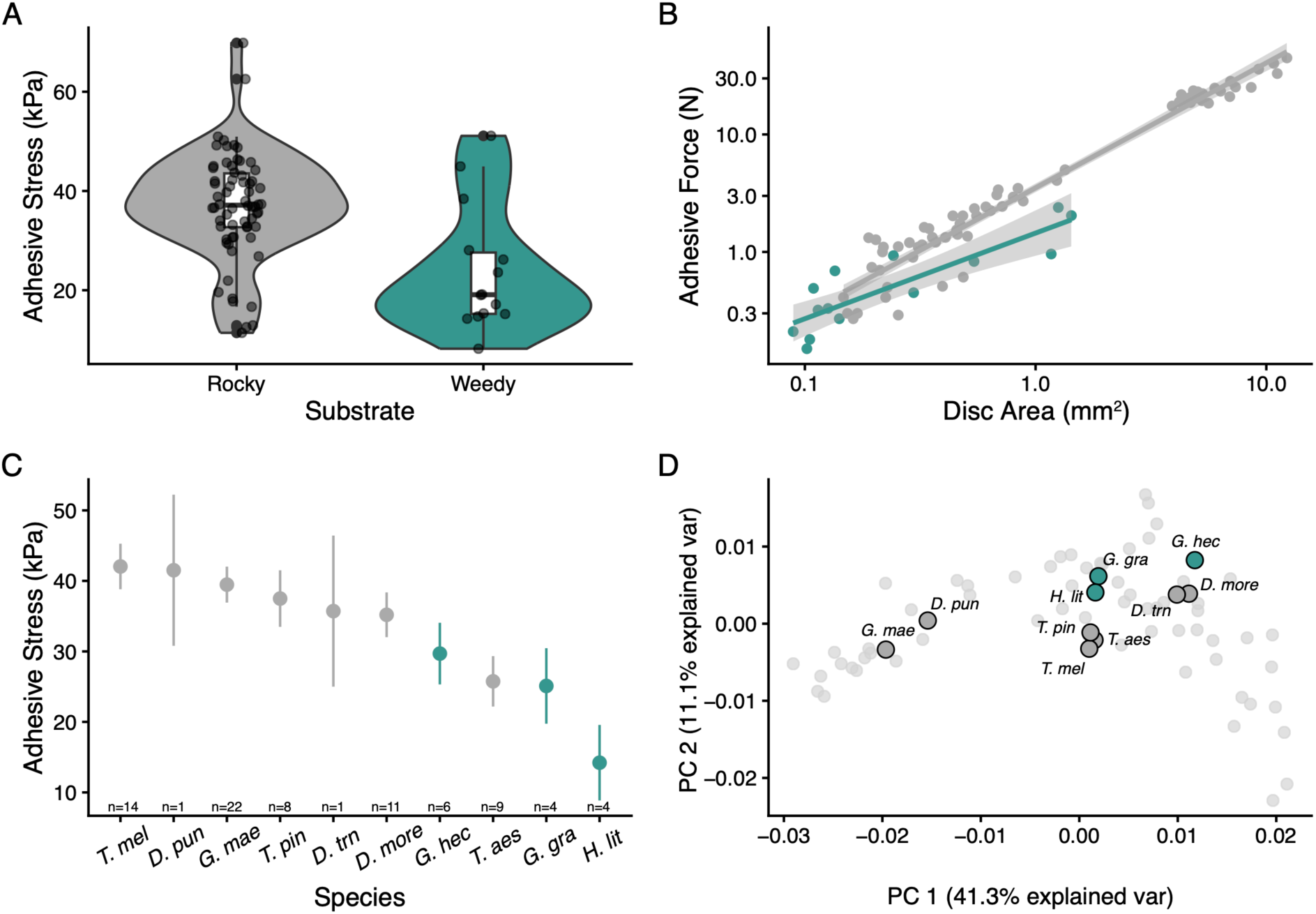
Variation in the adhesive performance of clingfishes. A) Comparison of max adhesive stress between rocky and weed-dwellers. B) Relationships between maximum suction force and disc area in rocky and weed-dwellers. C) Estimated marginal means and standard error bars showing interspecific variation in max adhesive stress. Also shown is the number of specimens sampled per species. D) Disc morphospace from Figure 3 highlighting the distribution of species that were sampled for performance.

Peak stress did not exhibit significant evolutionary integration with disc shape or either of the pectoral elements (p ≥ 0.061). However, peak stress did covary significantly with pelvic girdle shape (r-PLS = 0.93, p = 0.028). Greater adhesive stress was generally associated with wider and flatter pelvic girdles.

## DISCUSSION

Ventral adhesive discs are a hallmark of clingfish evolution and supports their diversification in benthic environments. In this study, we investigated the impact of habitat and substrate use on the clingfish skeleton. We found that habitat transitions to coral reefs promoted high levels of phenotypic disparity and faster rates of evolution, while the transition to freshwater had an opposite effect. We also found that weed-dwellers independently evolved at least six times and promoted morphological convergence in disc and skull shapes but constrained the overall disparity of disc shape. Differences in adhesive performance between rocky and a clade of weed-dwellers support a relationship between substrate use and disc morphology.

Overall, the skull evolved twice as fast as the disc, underscoring the diverse functions of the teleost skull and mosaic patterns of evolution across the clingfish body plan. Taken together, we propose that environmental demands and patterns of resource use are important factors shaping the evolutionary trajectories of clingfish morphology.

### Ecological covariates of adhesive disc form and function

We found strong evidence that substrate preferences impact the form and function of the adhesive discs. Convergence on a restricted region of the disc morphospace suggests that attaching to macroalgae and seagrass impose specific demands on clingfish discs. Specifically, the increased flexibility of weedy substrates may be more challenging to adhere to than hard rocky structures. This is evidenced by *G. maeandricus*, which generates lower adhesive force on more flexible substrates and biofouled surfaces compared to hard substrates (Ditsche et al. 2014; Huie and Summers, 2022). We posit that the disc shapes found in weed-dwellers may enhance attachment on softer substrates. The lower adhesive performance of weed-dwellers on hard substrates also suggests a potential trade-off between adhering to hard versus soft substrates.

However, we tested only a single origin of weed-dwellers (subfamily Haplocylicinae) and comparisons between more taxa and a wider range of substrates are needed to investigate these possible ecomechanical relationships. Alternatively, seagrass beds reduce wave energy (Fonseca and Cahalan, 1992) and may weaken the selective pressures on adhesive performance. This could explain why weed-dwellers have relatively smaller discs and generate less force per unit of disc area than rock-dwellers.

Surprisingly, the adhesive performance of rock-dwelling species seemed to vary minimally despite distinct disc skeletons. Maximum adhesive stress may be largely conserved and maintained by many-to-one mapping (Wainwright et al., 2005). If similar adhesive performance can be achieved through various morphological pathways, it could free up disc shape evolution to explore a range of designs. However, many-to-one mapping alone does not explain the patterns of disc diversity. We propose that variation in disc morphology reflects, in part, differences in microhabitat preferences and how much species rely on their discs to resist flow. For example, marine *Gobiesox* and *Gouania* are both intertidal species but *Gobiesox* are regularly battered with high-energy waves (Pires and Gibran, 2011; Distsche et al., 2017), while *Gouania* escape flow by burrowing in the sediment of gravel beaches (Wagner et al., 2019; 2023). Additionally, some clingfishes exhibit mixed substrate preferences that would also impact the selective pressures acting on their discs. *Cochloceps orientalis*, sets up cleaning stations on rocky and weedy substrates, and attach to their hosts as they clean (Hutchins, 1991). Others, such as *Apletodon*, exhibit ontogenetic variation and associate with algae and seagrass as juveniles and boulders as adults (Goncalves et al., 2002; Hofrichter and Patzner., 2001). Contrasting demands between rocky and weedy substrates could promote entirely different evolutionary trajectories compared to species with preferences for a singular substrate type.

Morphological variation in the clingfish disc may also be correlated with aspects of performance not measured in this study. For instance, live *G. maeandricus* can generate 1.3x more suction than dead ones, implying that muscles play an active role in modulating adhesion (Arnette et al., 2025). It is not known whether the performance of dead fishes is a strong predictor of live fish performance, but variation in disc musculature would likely contribute to differences in live adhesion. Major changes in pelvic girdle shape were associated with the relative size and arrangement of fenestra that are presumably correlated with muscle morphology (Arita, 1967). Specifically, flattening of the pelvic girdle and loss of the dorsal crest, as seen in *Gobiesox*, may accommodate relatively larger adductor muscles compared to smaller girdles with a crest that may restrict the space available for muscles. Disc muscles could also vary in the proportion of red versus white fibers, as seen in waterfall climbing gobies with different climbing strategies (Schoenfuss et al., 2013). Fiber types may be particularly relevant for managing fatigue in species that consistently engage their disc versus those that do so intermittently.

### Ecological axes of skull shape diversification

Consistent with the notion that coral reefs promote diversity (Alfaro et al., 2007; Kiesling et al., 2010; Price et al., 2011), we found that transitions to coral reef environments led to elevated rates of shape evolution across the clingfish skeleton. Reef species also explored novel regions of the two morphospaces investigated herein. Crinoid species, which are exclusive to coral reefs, exemplify how distinct reef species can be. *Discotrema* have novel disc components not found in any other clingfish, including a joint in the ventral postcleithra and hardened papillae on the ventral surface of the disc (Fujiwara et al., 2024) (Figure 5A). Meanwhile, *Rhinolepadichthys* feed on the pinnules of their crinoid hosts, expanding the trophic diversity of clingfishes (Fujiwara et al., 2024). These evolutionary shifts on coral reefs may reflect a release from the physical demands associated with harsh coastal environments, paired with the spectacular array of niches that reef systems are famous for. Indeed, reef species like *Diademichthys* or *Flabellicauda* have smaller adhesive discs and are the only epibenthic clingfishes, in that they are commonly observed hovering in the water column among sea urchin spines (Fujiwara et al., 2021). Taken together, it appears that clingfish have capitalized on opportunities for ecological and morphological specialization provided by coral reefs.

In contrast, freshwater transitions did not beget changes in skull shape or diversification. Marine-freshwater transitions can provide ecological opportunity and promote morphological diversity (Davis et al. 2012; Kolmann et al., 2020; de Brito et al., 2022), yet freshwater *Gobiesox* exhibited low levels of disparity and evolutionary rates. Competition with primary freshwater species is often used to explain the limited diversification of marine-derived freshwater lineages (Betancur-R et al., 2012). This could be the case for clingfishes, but we argue that it is not the only reason. Freshwater and marine *Gobiesox* were morphologically indistinguishable, which may reflect ecological conservatism (Buser et al., 2019). In freshwater systems, clingfish are found in fast flowing streams (Conway et al., 2017a) that likely impose similar hydrodynamic demands on head shape and adhesion as wave action in intertidal environments. Thus, the occupation of similar high-flow ecological niches in both marine and freshwater habitats is a plausible explanation for the lack of morphological diversification following a freshwater invasion.

Specialized microhabitats within the coastal environment also contributed to patterns of skull shape evolution. Weed-dwellers had convergent skulls that reflect adaptations for a cryptic lifestyle. Having a flatter and narrower skull allows weed-dwellers to more closely match the profile of a seagrass blade or a kelp stipe. Species that live on brown macroalgae or green seagrass also exhibit coloration that mimics their preferred substrates (Figure 1B, 1F), underscoring the importance of crypsis (Hofrichter and Patzner, 2000; Conway et al. 2024). The rapid skull evolution in *Alabes* was likely linked to their disc loss, but substrate preferences could help explain interspecific variation. *Alabes parvula* is a miniaturized and semi-transparent species, with possible developmental truncation (Britz and Conway, 2009; Springer and Fraser, 1976), that is associated with weedy structures while *A. dorsalis* occupies rockpools (Hutchins and Morrison, 2004). *Alabes dorsalis* has a similar skull shape and an eel-like body plan approaching that of the interstitial-dwelling *Gouania wildenowi*, suggesting that it may be adapted for moving through crevices between rocks, albeit above ground. In *Gouania,* modifications to the neurocrania associated with sediment composition were previously documented, whereby species living in interstices with smaller sediment grains have more slender neurocrania than those associated with larger sediment grains (Wagner et al., 2023; 2026).

Trophic ecology is another factor that often explains variation in jaws and dentitions but was not considered in this study. Many clingfish appear to feed on small invertebrates or zooplankton with some notable exceptions (Teytaud 1971; Critchlow 1972; Trkov et al., 2024). The largest clingfish species (*Chorisochismus dentex* and *Sicyayses sanguineus*) feed on larger and well-defended prey, such as limpets, mussels, and barnacles (Stobbs, 1980; Cancino and Castilla, 1988). The spatulate teeth of *S. sanguineus* and fangs of *C. dentex* may be adapted for prying prey off rocks (Conway et al., 2015) (Figure 5B). Furthermore, *Diademichthys* exhibit sexual divergence in diet and head shape, with females displaying a longer snout and stronger preference for shrimp eggs and bivalve gills than males, which feed on urchin tube feet (Sakashita, 1992). Other clingfishes have dimorphic head shapes, but they likely correlate more with male nest guarding and parental care than trophic partitioning (Conway et al., 2018). Thus, patterns of clingfish skull diversity are likely the result of complex interactions among habitat, diet, dimorphism and other unexamined factors.

### Future directions

We established links between ecology and skeletal morphology in clingfishes but the adhesive disc is comprised of important components that were not examined herein. Soft tissue traits, such as whole disc shape, papillae morphology, and musculature, require further investigation to better understand the relationships between disc form and function. For example, adhesive discs vary in surface complexity and comprise either a single complete circular structure or an incomplete arc with a small, inner disc (Briggs, 1955). How these different shapes influence adhesion is not known, but bioinspired suction cups indicate that disc shape affects how well discs resist shear forces (Hernandez et al., 2024). Related to this, the papillae on the ventral surface of clingfish adhesive discs vary in their size, density, and arrangement across species, suggesting that they also vary with environmental demands (Sandoval et al., 2020; Hernandez et al., 2025). Alternatively, the papillae biofluoresce in some species and might serve roles unrelated to adhesion, such as communication (Cohen et al., 2021; Huie et al., 2022; Carr et al., 2025). Follow up experiments that investigate the performance of disc shapes are needed, such as ones that compare the performance of live versus euthanized animals or investigate trade-offs between adhering to soft and hard substrates. These endeavors will provide a deeper understanding of the ecological and evolutionary mechanisms shaping disc morphology and may provide source material for advancing the development of bioinspired technology.

## AUTHOR CONTRIBUTIONS

JMH - Conceptualization, Data curation, Formal analysis, Investigation, Methodology, Visualization, Writing—original draft; APS – Conceptualization, Data curation, Funding acquisition, Investigation, Methodology, Resources, Writing – review & editing; KCH – Investigation, Methodology, Writing – review & editing; TT - Investigation, Resources, Writing – review & editing; SH - Investigation, Resources, Writing – review & editing; GV - Methodology, Writing – review & editing; CMM - Methodology, Writing – review & editing; KWC – Conceptualization, Investigation, Methodology, Resources, Writing – review & editing

## FUNDING

This research was supported by the National Science Foundation (DGE-1746914 to J.M.H, DBI-1701665 and IOS-1256602 to A.P.S., DBI-1702442 and IOS-1256793 to K.W.C.) and Texas A&M Agrilife Research (Hatch TEX09452-1 to K.W.C.).

## CONFLICT OF INTERESTS

The authors declare that they have no known competing financial interests or personal relationships that could have appeared to influence the work reported in this paper.

## ACKNOWLEDGEMENTS

We thank the many natural history collections and their staff for providing access to their specimens for imaging. We also thank Andrew Williston (MCZ) for CT scanning *Apletodon incognitus* and James Maclaine, Brett Clark, and Vincent Fernandez for scanning *Opeatogenys gracilis* for this study. We thank Richard Blob for providing a scan of *Gobiesox cephalus* and Karly Cohen for their assistance with scanning. Additionally, we thank Kory Evans for their helpful conversations on fish skulls and for providing analysis scripts. We also thank Barry Brown/coralreefphotos, Graham Short, Ian Sipworth, Mark Erdmann, Maximilian Wagner, and Rudie Kuiter for allowing us to use their clingfish photographs.

## Notes

### Competing Interest Statement

The authors have declared no competing interest.

### Summary of Updates

Corrected a spelling error with co-author's name and updated funding information.

