## Supplemental Information for "Habitat impacts the diversification of adhesive discs and skull shape in clingfishes"

### In this document:

Supplemental Tables
- Table S1: List of 82 clingfishes sampled in the phylogeny and ecological classifications.
- Table S2: List of 74 clingfishes sampled for morphology and specimen metadata.
- Table S3: List of landmarks used for 3D geometrics morphometrics.
- Table S4: Maximum adhesive stress for 10 species of clingfishes.
- Table S5: Results of the MANOVAs testing the effects of ecology on shape.

Supplemental Figures
- Figure S1: Ancestral state reconstruction of habitat and substrate use.
- Figure S2: Representation of the landmarks placed on each of the bones.

Supplemental References

**Table S1.** The 82 extant clingfish species included in the phylogeny and their habitat and substrate classifications.

| Species | Subfamily | Habitat | Substrate | Citations |
|---|---|---|---|---|
| *Acyrtops beryllinus* | Gobiesocinae | Coastal | Weedy | Gould, 1965 |
| *Acyrtus artius* | Gobiesocinae | Coral Reef | Rocky | Johnson & Greenfield, 1983 |
| *Acyrtus lanthanum* | Gobiesocinae | Coral Reef | Rocky | Conway et al., 2014 |
| *Acyrtus rubiginosus* | Gobiesocinae | Coral Reef | Rocky | Johnson & Greenfield, 1983 |
| *Alabes dorsalis* | Cheilobranchinae | Coastal | Rocky | Kuiter, 1993; Hutchins & Morrison, 2004 |
| *Alabes parvula* | Cheilobranchinae | Coastal | Weedy | Kuiter, 1993; Hutchins & Morrison, 2004 |
| *Apletodon dentatus* | Lepadogastrinae | Coastal | Rocky | Gonçalves et al., 2002 |
| *Apletodon incognitus* | Lepadogastrinae | Coastal | Rocky | Patzner, 1999; Hofrichter & Patzner, 2000 |
| *Arcos erythrops* | Gobiesocinae | Coastal | Rocky | FishBase |
| *Aspasma ubauo* | Diademichthyinae | Coastal | Weedy | Masuda et al., 1984; Fujiwara & Motomura, 2020 |
| *Aspasmichthys ciconiae* | Diademichthyinae | Coastal | Rocky | Masuda et al., 1984 |
| *Aspasmogaster costata* | *Incertae sedis* | Coastal | Rocky | FishBase |
| *Aspasmogaster liorhynchus* | *Incertae sedis* | Coastal | Rocky | FishBase |
| *Aspasmogaster tasmaniensis* | *Incertae sedis* | Coastal | Rocky | FishBase |
| *Barryichthys hutchinsi* | Cheilobranchinae | Coastal | Weedy | Conway et al., 2018 |
| *Chorisochismus dentex* | Chorisochisminae | Coastal | Rocky | Stobbs, 1980 |
| *Cochleoceps bassensis* | Cheilobranchinae | Coastal | Rocky | Hutchins, 2008 |
| *Cochleoceps orientalis* | Cheilobranchinae | Coastal | Rocky | Kuiter, 1993 |
| *Cochleoceps spatula* | Cheilobranchinae | Coastal | Weedy | Hyndes et al., 2003 |
| *Cochleoceps viridis* | Cheilobranchinae | Coastal | Weedy | Hyndes et al., 2003 |
| *Conidens laticephalus* | *Incertae sedis* | Coastal | Rocky | Shiogaki & Dotsu, 1971; Masuda et al., 1984 |
| *Creocele cardinalis* | *Incertae sedis* | Coastal | Rocky | Kuiter, 1993 |
| *Dellichthys morelandi* | Trachelochisminae | Coastal | Rocky | Stewart, 2015 |
| *Dellichthys trnskii* | Trachelochisminae | Coastal | Rocky | Conway et al., 2018 |
| *Derilissus nanus* | Gobiesocinae | Coral Reef | Rocky | Sparks & Gruber, 2012 |
| *Diademichthys lineatus* | Diademichthyinae | Coral Reef | Rocky | FishBase |
| *Diplecogaster bimaculata* | Lepadogastrinae | Coastal | Rocky | Hofrichter & Patzner, 2001 |
| *Diplocrepis puniceus* | Diplocrepinae | Coastal | Rocky | FishBase; Stewart, 2015 |
| *Discotrema crinophilum* | Diademichthyinae | Coral Reef | Crinoid | Briggs, 1976 |
| *Eckloniaichthys scylliorhiniceps* | Chorisochisminae | Coastal | Weedy | Allen & Griffiths, 1981 |
| *Erdmannichthys alorensis* | Diademichthyinae | Coral Reef | Rocky | Allen & Erdmann, 2012; Conway et al., 2021 |
| *Flabellicauda akiko* | Diademichthyinae | Coral Reef | Rocky | Allen & Erdmann, 2012; Fujiwara et al., 2021 |
| *Flabellicauda alleni* | Diademichthyinae | Coral Reef | Rocky | Fujiwara et al., 2021 |
| *Flexor incus* | Diademichthyinae | Coastal | Rocky | Stewart, 2015; Conway et al., 2018 |
| *Gastrocyathus gracilis* | Haplocylicinae | Coastal | Weedy | Stewart, 2015 |
| *Gastrocymba quadriradiata* | Haplocylicinae | Coastal | Weedy | Stewart, 2015 |
| *Gastroscyphus hectoris* | Haplocylicinae | Coastal | Weedy | Stewart, 2015 |
| *GenusA sp1* | Cheilobranchinae | Coastal | Weedy | Hutchins, 1994; 2008 |
| *Gobiesox adustus* | Gobiesocinae | Coastal | Rocky | Castellanos-Galindo et al., 2005 |
| *Gobiesox barbatulus* | Gobiesocinae | Coastal | Rocky | Johnson & Greenfield, 1983; Pires & Gibran, 2011 |
| *Gobiesox cephalus* | Gobiesocinae | Freshwater | Rocky | Forks et al., 2014 |
| *Gobiesox daedalus* | Gobiesocinae | Coastal | Rocky | Gonzalez-Murcia et al., 2016 |
| *Gobiesox funebris* | Gobiesocinae | Coastal | Rocky | Eger, 1971 |
| *Gobiesox juradoensis* | Gobiesocinae | Freshwater | Rocky | Fowler, 1944 |
| *Gobiesox maeandricus* | Gobiesocinae | Coastal | Rocky | Johnson, 1970 |
| *Gobiesox mexicanus* | Gobiesocinae | Freshwater | Rocky | Mercado-Silva et al., 2016 |
| *Gobiesox nigripinnis* | Gobiesocinae | Coastal | Rocky | Conway et al., 2017 |
| *Gobiesox pinniger* | Gobiesocinae | Coastal | Rocky | FishBase |
| *Gobiesox potamius* | Gobiesocinae | Freshwater | Rocky | FishBase |
| *Gobiesox punctulatus* | Gobiesocinae | Coastal | Rocky | Johnson & Greenfield, 1983 |
| *Gobiesox rhessodon* | Gobiesocinae | Coastal | Rocky | FishBase |

| | | | | |
|---|---|---|---|---|
| *Gobiesox strumosus* | Gobiesocinae | Coastal | Rocky | FishBase |
| *Gouania willdenowi* | Lepadogastrinae | Coastal | Rocky | Patzner, 1998; Hofritcher & Patzner, 2000 |
| *Haplocylix littoreus* | Haplocylicinae | Coastal | Weedy | Stewart, 2015 |
| *Kopua minima* | Protogobiesocinae | Deep | Rocky | Moore et al., 2012 |
| *Lepadichthys frenatus* | Diademichthyinae | Coastal | Rocky | Masuda et al., 1984; Fujiwara & Motomura, 2019 |
| *Lepadichthys trishula* | Diademichthyinae | Coral Reef | Rocky | Fujiwara et al., 2020 |
| *Lepadogaster candollei* | Lepadogastrinae | Coastal | Rocky | Gonçalves et al. 1998; Hofritcher & Patzner, 2000 |
| *Lepadogaster lepadogaster* | Lepadogastrinae | Coastal | Rocky | Hofritcher & Patzner, 2000 |
| *Lepadogaster purpurea* | Lepadogastrinae | Coastal | Rocky | Gonçalves et al., 1998 |
| *Melanophorichthys posidonus* | Cheilobranchinae | Coastal | Weedy | Conway et al., 2024 |
| *Opeatogenys gracilis* | Lepadogastrinae | Coastal | Weedy | Hofritcher & Patzner, 2000 |
| *Parvicrepis parvipinnis* | Cheilobranchinae | Coastal | Weedy | Kuiter, 1993 |
| *Parvicrepis sp* | Cheilobranchinae | Coastal | Weedy | Kuiter, 1993 |
| *Pherallodichthys sp* | Diademichthyinae | Coral Reef | Rocky | FishBase |
| *Pherallodus indicus* | Diademichthyinae | Coastal | Rocky | Masuda et al., 1984 |
| *Posidonichthys hutchinsi* | Cheilobranchinae | Coastal | Weedy | FishBase |
| *Protogobiesox asymmetricus* | Protogobiesocinae | Deep | Rocky | Fricke et al. 2017 |
| *Rhinolepadichthys polyastrous* | Diademichthyinae | Coral Reef | Crinoid | Fujiwara et al., 2024 |
| *Rimicola muscarum* | Gobiesocinae | Coastal | Weedy | Rolan, 1978 |
| *Sicyases sanguineus* | Gobiesocinae | Coastal | Rocky | Cancino & Castillo, 1998 |
| *Tomicodon boehlkei* | Gobiesocinae | Coastal | Rocky | Eger, 1971; Critchlow, 1972 |
| *Tomicodon briggsi* | Gobiesocinae | Coastal | Rocky | Williams & Tyler, 2003 |
| *Tomicodon fasciatus* | Gobiesocinae | Coastal | Rocky | Johnson & Greenfield, 1983 |
| *Tomicodon humeralis* | Gobiesocinae | Coastal | Rocky | Eger, 1971 |
| *Tomicodon lavettsmithi* | Gobiesocinae | Coastal | Rocky | Williams & Tyler, 2003 |
| *Tomicodon myersi* | Gobiesocinae | Coastal | Rocky | Critchlow, 1972 |
| *Tomicodon reitzae* | Gobiesocinae | Coastal | Rocky | Williams & Tyler, 2003 |
| *Tomicodon zebra* | Gobiesocinae | Coastal | Rocky | Critchlow, 1972 |
| *Trachelochismus aestuarium* | Trachelochisminae | Coastal | Rocky | Conway et al., 2017 |
| *Trachelochismus melobesia* | Trachelochisminae | Coastal | Rocky | Stewart, 2015 |
| *Trachelochismus pinnulatus* | Trachelochisminae | Coastal | Rocky | Stewart, 2015 |

**Table S2.** The 74 extant clingfish species sampled for morphology, their catalogue numbers, and link to CT dataset on MorphoSource.

| Species | Museum | MorphoSource |
|---|---|---|
| *Acyrtops beryllinus* | TCWC 16774.01 | https://www.morphosource.org/media/000030729 |
| *Acyrtus artius* | ANSP 81299 | https://www.morphosource.org/media/000030726 |
| *Acyrtus lanthanum* | UF 212697 | https://www.morphosource.org/media/000899059 |
| *Acyrtus rubiginosus* | UF 149202 | https://www.morphosource.org/media/000030730 |
| *Alabes dorsalis* | AMNH 48737 | https://www.morphosource.org/media/000899065 |
| *Alabes parvula* | TCWC 17165.05 | https://www.morphosource.org/media/000030744 |
| *Apletodon incognitus* | MCZ 12940 | https://www.morphosource.org/media/000163335 |
| *Arcos erythrops* | FMNH 62067 | https://www.morphosource.org/media/000030763 |
| *Aspasma ubauo* | NSMT-P 114699 | https://www.morphosource.org/media/000899078 |
| *Aspasmichthys ciconiae* | TCWC 16461.02 | https://www.morphosource.org/media/000030899 |
| *Aspasmogaster costata* | TCWC 17166.01 | https://www.morphosource.org/media/000030754 |
| *Aspasmogaster liorhynchus* | NMV A 2558 | https://www.morphosource.org/media/000030831 |
| *Aspasmogaster tasmaniensis* | NMV A 3000 | https://www.morphosource.org/media/000030760 |
| *Barryichthys hutchinsi* | AMS I.49000-001 | https://www.morphosource.org/media/000080016 |
| *Chorisochismus dentex* | ROM 50997 | https://www.morphosource.org/media/000013792 |
| *Cochleoceps bassensis* | NMV A 8875 | https://www.morphosource.org/media/000030761 |
| *Cochleoceps orientalis* | AMS I.43845-005 | https://www.morphosource.org/media/000899584 |
| *Cochleoceps spatula* | WAM P.28288-002 | https://www.morphosource.org/media/000031251 |
| *Cochleoceps viridis* | WAM P.30262-001 | https://www.morphosource.org/media/000031252 |
| *Conidens laticephalus* | NSMT-P 58073 | https://www.morphosource.org/media/000899609 |
| *Creocele cardinalis* | NMV A 20189-001 | https://www.morphosource.org/media/000021573 |
| *Dellichthys morelandi* | NMNZ P.030626 | https://www.morphosource.org/media/000037807 |
| *Dellichthys trnskii* | NMNZ P.028060 | https://www.morphosource.org/media/000040584 |
| *Derilissus nanus* | UF 13372 | https://www.morphosource.org/media/000030758 |
| *Diademichthys lineatus* | ROM 74261 | https://www.morphosource.org/media/000899613 |
| *Diplecogaster bimaculata* | ROM 41585 | https://www.morphosource.org/media/000030901 |
| *Diplocrepis puniceus* | NMNZ P037447 | https://www.morphosource.org/media/000899618 |
| *Discotrema crinophilum* | ROM 85350 | https://www.morphosource.org/media/000030750 |
| *Eckloniaichthys scylliorhiniceps* | SAIAB 17-083 | https://www.morphosource.org/media/000899624 |
| *Erdmannichthys alorensis* | WAM P.34629-005 | https://www.morphosource.org/media/000388066 |
| *Flabellicauda akiko* | WAM P.35180-001 | https://www.morphosource.org/media/000899646 |
| *Flabellicauda alleni* | ROM 55185 | https://www.morphosource.org/media/000030731 |
| *Flexor incus* | AIM MA655316 | https://www.morphosource.org/media/000056344 |
| *Gastrocyathus gracilis* | AIM MA6871 | https://www.morphosource.org/media/000899629 |
| *Gastrocymba quadriradiata* | NMNZ P.053921 | https://www.morphosource.org/media/000899635 |
| *Gastroscyphus hectoris* | AIM MA130487 | https://www.morphosource.org/media/000899640 |
| *GenusA sp1* | NMV A-20781 | https://www.morphosource.org/media/000899651 |
| *Gobiesox adustus* | LACM 1593 | https://www.morphosource.org/media/000655614 |
| *Gobiesox cephalus* | Blob unCat | https://www.morphosource.org/media/000902519 |
| *Gobiesox daedalus* | UWFC 22587 | https://www.morphosource.org/media/000899675 |
| *Gobiesox funebris* | SIO 87-178 | https://www.morphosource.org/media/000030765 |

| | | |
|---|---|---|
| *Gobiesox juradoensis* | USNM 360154 | https://www.morphosource.org/media/000052687 |
| *Gobiesox maeandricus* | UWFC 000643 | https://www.morphosource.org/media/000078707 |
| *Gobiesox mexicanus* | UMMZ 223307 | https://www.morphosource.org/media/000902468 |
| *Gobiesox nigripinnis* | UF 149209 | https://www.morphosource.org/media/000052678 |
| *Gobiesox pinniger* | SIO 65-286 | https://www.morphosource.org/media/000030822 |
| *Gobiesox potamius* | NRM 53428 | https://www.morphosource.org/media/000902480 |
| *Gobiesox punctulatus* | TCWC 16451.01 | https://www.morphosource.org/media/000902474 |
| *Gobiesox rhessodon* | UF 10169 | https://www.morphosource.org/media/000902493 |
| *Gobiesox strumosus* | TCWC 11200.20 | https://www.morphosource.org/media/000030764 |
| *Gouania willdenowi* | TCWC 16777.03 | https://www.morphosource.org/media/000027515 |
| *Haplocylix littoreus* | UF 80190 | https://www.morphosource.org/media/000051236 |
| *Lepadichthys frenatus* | ROM 65285 | https://www.morphosource.org/media/000029847 |
| *Lepadogaster candollei* | UF 209719 | https://www.morphosource.org/media/000902500 |
| *Lepadogaster lepadogaster* | UF 81193 | https://www.morphosource.org/media/000902512 |
| *Lepadogaster purpurea* | NMBE 1078409 | https://www.morphosource.org/media/000903818 |
| *Melanophorichthys posidonus* | AMS I.51546-001 | https://www.morphosource.org/media/000665747 |
| *Opeatogenys gracilis* | NHM 1995.3.15.1 | https://www.morphosource.org/media/000902698 |
| *Parvicrepis parvipinnis* | TCWC 17169.01 | https://www.morphosource.org/media/000030755 |
| *Parvicrepis sp* | TCWC 17169.02 | https://www.morphosource.org/media/000902655 |
| *Pherallodichthys sp* | ROM 55170 | https://www.morphosource.org/media/000902671 |
| *Pherallodus indicus* | NSMT-P 114703 | https://www.morphosource.org/media/000056337 |
| *Posidonichthys hutchinsi* | KWC 16-01 | https://www.morphosource.org/media/000030706 |
| *Rhinolepadichthys polyastrous* | ROM 72940 | https://www.morphosource.org/media/000029846 |
| *Rimicola muscarum* | ANSP 92371 | https://www.morphosource.org/media/000031667 |
| *Sicyases sanguineus* | AMNH 37963 | https://www.morphosource.org/media/000903781 |
| *Tomicodon boehlkei* | UF 26496 | https://www.morphosource.org/media/000902709 |
| *Tomicodon fasciatus* | ANSP 81295 | https://www.morphosource.org/media/000030906 |
| *Tomicodon humeralis* | LACM 30126.001 | https://www.morphosource.org/media/000655859 |
| *Tomicodon myersi* | LACM 51360.003 | https://www.morphosource.org/media/000656083 |
| *Tomicodon zebra* | MCZ 45664 | https://www.morphosource.org/media/000030900 |
| *Trachelochismus aestuarium* | TCWC 17266.03 | https://www.morphosource.org/media/000902843 |
| *Trachelochismus melobesia* | TCWC 17174.01 | https://www.morphosource.org/media/000030694 |
| *Trachelochismus pinnulatus* | AIM MA5220 | https://www.morphosource.org/media/000903243 |

**Table S3.** List of fixed and semi-landmark curves used for 3D geometric morphometrics.

| Landmark # | Bone | Description |
|---|---|---|
| 1 | Premaxilla | Most distal point of first tooth on premaxilla |
| 2 | Premaxilla | Base of first premaxilla tooth |
| 3 | Premaxilla | Most distal point of ascending process of premaxilla |
| 4 | Premaxilla | Proximal vertex between ascending and descending processes of premaxilla |
| 5 | Premaxilla | Most distal point of descending process of premaxilla |
| 6 | Dentary | Most distal point of first tooth on dentary |
| 7 | Dentary | Posterior-most point on dentary flange |
| 8 | Dentary | Ventral-most point of mental symphysis on dentary |
| 9 | Angular-Articular | Antero-distal-most point of articular/angular |
| 10 | Angular-Articular | Distal-most point of coronoid process |
| 11 | Dentary | Ventral, posterior-most point of dentary |
| 12 | Angular-Articular | Center of jaw joint on quadrate |
| 13 | Angular-Articular | Posterior-most point on retroarticular |
| 14 | Neurocranium | Most anterior point of vomer |
| 15 | Neurocranium | Left, distal-most point on anterior of vomer |
| 16 | Neurocranium | Distal-most point of the lateral ethmoid |
| 17 | Neurocranium | Proximal lateral ethmoid-parasphenoid margin |
| 18 | Neurocranium | Dorsal, medial-most point of the ethmoid |
| 19 | Neurocranium | Proximal-most point of lateral ethmoid frontal margin |
| 20 | Neurocranium | Origin of supraoccipital |
| 21 | Neurocranium | Posterior frontal sub-orbital margin |
| 22 | Neurocranium | Pterosphenoid-parasphenoid margin |
| 23 | Neurocranium | Mid-point Parasphenoid-basioccipital margin |
| 24 | Neurocranium | Medial edge of the prootic foramen |
| 25 | Neurocranium | Posterior-most point of supraoccipital |
| 26 | Neurocranium | Ventral epiotic-post temporal margin |
| 27 | Neurocranium | Distal-most dorsal point of basioccipital |
| 28 | Neurocranium | Pterotic-post temporal margin |
| 29 | Neurocranium | Distal-most ventral point of basioccipital |
| 30-39 | Premaxilla | Curve 1: Ascending process of premaxilla, between landmarks 2 and 3 |
| 41-44 | Dentary | Curve 2: Ventral margin of dentary, between landmarks 8 and 11 |
| 45-54 | Neurocranium | Curve 3: Ventral margin of parasphenoid, between landmarks 14 and 23 |
| 55-63 | Neurocranium | Curve 4: Orbital margin, between landmarks 18 and 21 |
| 64 | Maxilla | Distal, dorsal anterior margin of maxilla |
| 65 | Maxilla | Distal, dorsal posterior margin of maxilla |
| 66 | Maxilla | Distal, ventral anterior margin of maxilla |
| 67 | Maxilla | Distal, ventral posterior margin of maxilla |
| 68 | Maxilla | Proximal,dorsal anterior margin of maxilla |
| 69 | Maxilla | Proximal,dorsal posterior margin of maxilla |
| 70 | Maxilla | Anterior-most point of the proximal descending process of maxilla |
| 71 | Hyomandibula | Anterior-most pont of contact on the distal face of the hyomandibula between hyomandibula and sphenotic |

| | | |
|---|---|---|
| 72 | Hyomandibula | Posterior-most point of contact on the distal face of the hyomandibula between hyomandibula and the pterotic |
| 73 | Hyomandibula | Distal-most point of lateral projection of hyomandibula |
| 74 | Hyomandibula | Anterior-ventral most point of hyomandibula |
| 75 | Hyomandibula | Ventral-most point of hyomandibula |
| 76-79 | Angular-Articular | Curve 5: Angular ascending process, between landmarks 10 and 12 |
| 80-83 | Angular-Articular | Curve 6: Angular lateral process, between landmarks 9 and 12 |
| 84-88 | Premaxilla | Curve 7: Lateral arm of premaxilla, between landmarks 2 and 5 |
| 89-92 | Hyomandibula | Curve 8: Anterior face of hyomandibula, between landmarks 73 and 74 |
| 93-96 | Hyomandibula | Curve 9: Dorsal surface of hyomandibula, between landmarks 71 and 72 |
| 97-100 | Hyomandibula | Curve 10: Posterior face of hyomandibula, between landmarks 73 and 75 |
| 101 | Urohyal | Dorsal, anterior-most point of urohyal |
| 102 | Urohyal | Ventral anterior-most point of urohyal |
| 103 | Urohyal | Dorsal, posterior-most point of urohyal |
| 104 | Urohyal | Ventral, posterior-most point of urohyal |
| 105 | Hyoid | Anterior-most point of ceratohyal |
| 106 | Hyoid | Interior-ridge of ceratohyal |
| 107 | Hyoid | Dorsal ceratohyal-epihyal margin |
| 108 | Hyoid | Posterior-most point of epihyal |
| 109 | Hyoid | Ventral ceratohyal-epihyal margin |
| 110-114 | Urohyal | Curve 11: Urohyal dorsal edge, between landmarks 101 and 103 |
| 115-119 | Urohyal | Curve 12: Urohyal ventral edge, between landmarks 102 and 104 |
| 120-124 | Hyoid | Curve 13: Ceratohyal dorsal ridge, between landmarks 105 and 107 |
| 125-129 | Hyoid | Curve 14: Ceratohyal ventral ridge, between landmarks 105 and 109 |
| 130-134 | Neurocranium | Curve 15: Supraoccipital crest, between landmarks 20 and 25 |
| 135 | Pelvic Girdle | Posterior-most point of the basipterygium |
| 136 | Pelvic Girdle | Posterior-most point of articulation with the fifth pelvic fin ray |
| 137 | Pelvic Girdle | Posterior margin of the articulation between the first pelvic fin ray and basipterygium |
| 138 | Pelvic Girdle | Posterior point of contact between pectoral girdle and basipterygium |
| 139 | Pelvic Girdle | Anterior margin of the internal process along the basipterygium midline |
| 140 | Pelvic Girdle | Posterior margin of the internal process along the basipterygium midline |
| 141 | Pelvic Girdle | Anterior-ventral most point along the basipterygium midline |
| 142 | Pelvic Girdle | On the ventral surface of the basipterygium, the posterior-medial margin of the anterolateral fenestrum |
| 143 | Pelvic Girdle | On the ventral surface of the basipterygium, the anterior-lateral margin of the central fenestrum |
| 144 | Pelvic Girdle | Anterior-most point of the external process |
| 145 | Pelvic Girdle | On the ventral surface of the basipterygium, the posterior-medial margin of the central fenestrum |
| 146 | Pelvic Girdle | On the dorsal surface of the basipterygium, the lateral margin of the central fenestrum |
| 147-152 | Pelvic Girdle | Curve 16: Basipterygium dorsal ridge, between landmarks 135 and 140 |
| 153-157 | Pelvic Girdle | Curve 17: Basipterygium lateral ridge, between landmarks 135 and 136 |
| 158-162 | Pelvic Girdle | Curve 18: Basipterygium ventral midline, between landmarks 135 and 141 |
| 163 | Pectoral Girdle | Dorsal-most point of pectoral symphysis |
| 164 | Pectoral Girdle | Ventral-most point of pectoral symphysis |

| | | |
|---|---|---|
| 165 | Pectoral Girdle | Anterior margin of the pectoral girlde and basipterygium joint |
| 166 | Pectoral Girdle | Posterior margin of the pectoral girlde and basipterygium joint |
| 167 | Pectoral Girdle | Dorsal-most point on the lateral margin of the cleithrum |
| 168 | Pectoral Girdle | Dorsal-margin of the articulation with the supracleithrum |
| 169 | Pectoral Girdle | Inflection point along the medial margin of the cleithrum |
| 170 | Pectoral Girdle | Medial and dorsal-most point of the upper arm of the cleithrum |
| 171 | Pectoral Girdle | Lateral and dorsal-most point of the upper arm of the cleithrum |
| 172-177 | Pectoral Girdle | Curve 19: Cleithrum lateral edge, between landmarks 163 and 167 |
| 178 | Ventral Postcleithrum | Anterior-medial edge of articulating surface with basipterygium (in some) |
| 179-195 | Ventral Postcleithrum | Curve 20: Ventral postcleithrum perimeter, start and end at landmark 178. Clockwise from dorsal view |

**Table S4.** The estimated marginal mean and standard error of maximum adhesive stress (kPa) for 10 species of clingfishes tested on a substrate with surface roughness of 15.3 µm.

| Species | Subfamily | n | emmean | SE |
|---|---|---|---|---|
| *Dellichthys morelandi* | Trachelochisminae | 14 | 35.2 | 3.17 |
| *Dellichthys trnskii* | Trachelochisminae | 1 | 35.7 | 10.7 |
| *Diplocrepis puniceus* | Diplocrepinae | 1 | 41.5 | 10.7 |
| *Gastrocyathus gracilis* | Haplocylicinae | 4 | 25.1 | 5.36 |
| *Gastroscyphus hectoris* | Haplocylicinae | 6 | 29.7 | 4.38 |
| *Gobiesox maeandricus* | Gobiesocinae | 22 | 39.5 | 2.55 |
| *Haplocylix littoreus* | Haplocylicinae | 4 | 14.2 | 5.36 |
| *Trachelochismus aestuarium* | Trachelochisminae | 9 | 25.8 | 3.57 |
| *Trachelochismus melobesia* | Trachelochisminae | 11 | 42.0 | 3.23 |
| *Trachelochismus pinnulatus* | Trachelochisminae | 8 | 37.5 | 4.01 |

**Table S5.** Phylogenetic MANOVA results testing the effects of habitat and substrate on disc and skull shape.

| Trait | Fixed Effects | Df | SS | MS | Rsq | F | Z | p-value |
|---|---|---|---|---|---|---|---|---|
| Disc Shape | Size | 1 | 6.97E-05 | 6.97E-05 | 0.009 | 0.644 | 0.570 | 0.297 |
| | Habitat | 2 | 6.69E-05 | 3.34E-05 | 0.009 | 0.309 | -0.950 | 0.819 |
| | Size * Habitat | 2 | 4.60E-04 | 2.30E-04 | 0.059 | 2.123 | 0.828 | 0.211 |
| | Size | 1 | 9.13E-05 | 9.13E-05 | 0.012 | 0.832 | 0.898 | 0.213 |
| | Substrate | 2 | 2.00E-04 | 1.00E-04 | 0.026 | 0.911 | 0.323 | 0.406 |
| | Size * Substrate | 2 | 2.06E-04 | 1.03E-04 | 0.027 | 0.938 | 0.366 | 0.367 |
| Skull Shape | Size | 1 | 8.40E-05 | 8.40E-05 | 0.024 | 1.792 | 1.394 | 0.086 |
| | Habitat | 2 | 7.99E-05 | 4.00E-05 | 0.023 | 0.852 | 0.448 | 0.335 |
| | Size * Habitat | 2 | 1.19E-04 | 5.93E-05 | 0.034 | 1.264 | 0.602 | 0.27 |
| | Size | 1 | 8.40E-05 | 8.40E-05 | 0.024 | 1.772 | 1.383 | 0.092 |
| | Substrate | 2 | 6.43E-05 | 3.21E-05 | 0.019 | 0.678 | 0.171 | 0.427 |
| | Size * Substrate | 2 | 9.86E-05 | 4.93E-05 | 0.028 | 1.040 | 0.383 | 0.354 |

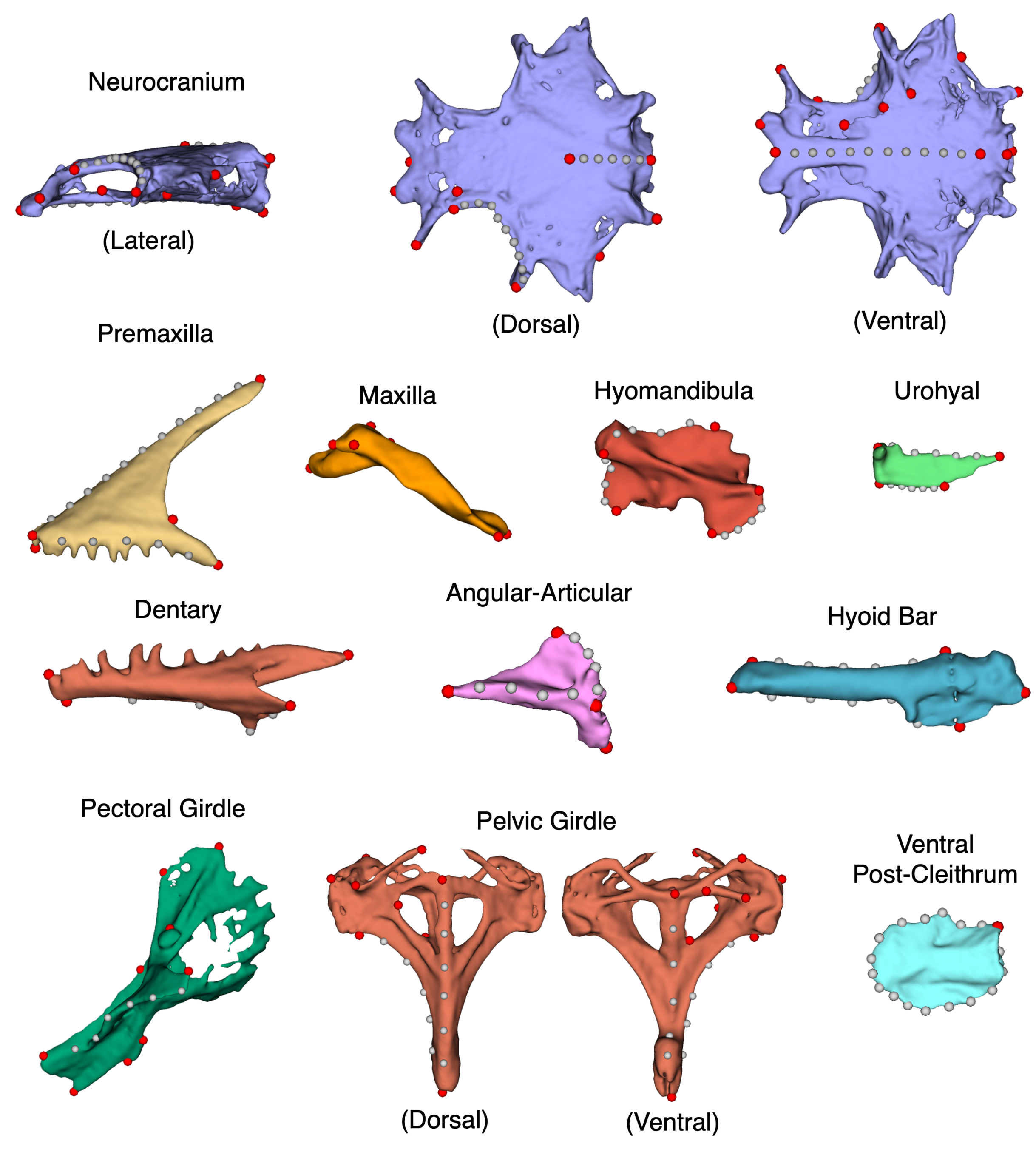


**Figure S1.** Representation of the landmarks placed on the different bones for *Conidens laticephalus* (NSMT-P 58073). Fixed landmarks are in red, semi-landmark curves are in grey.

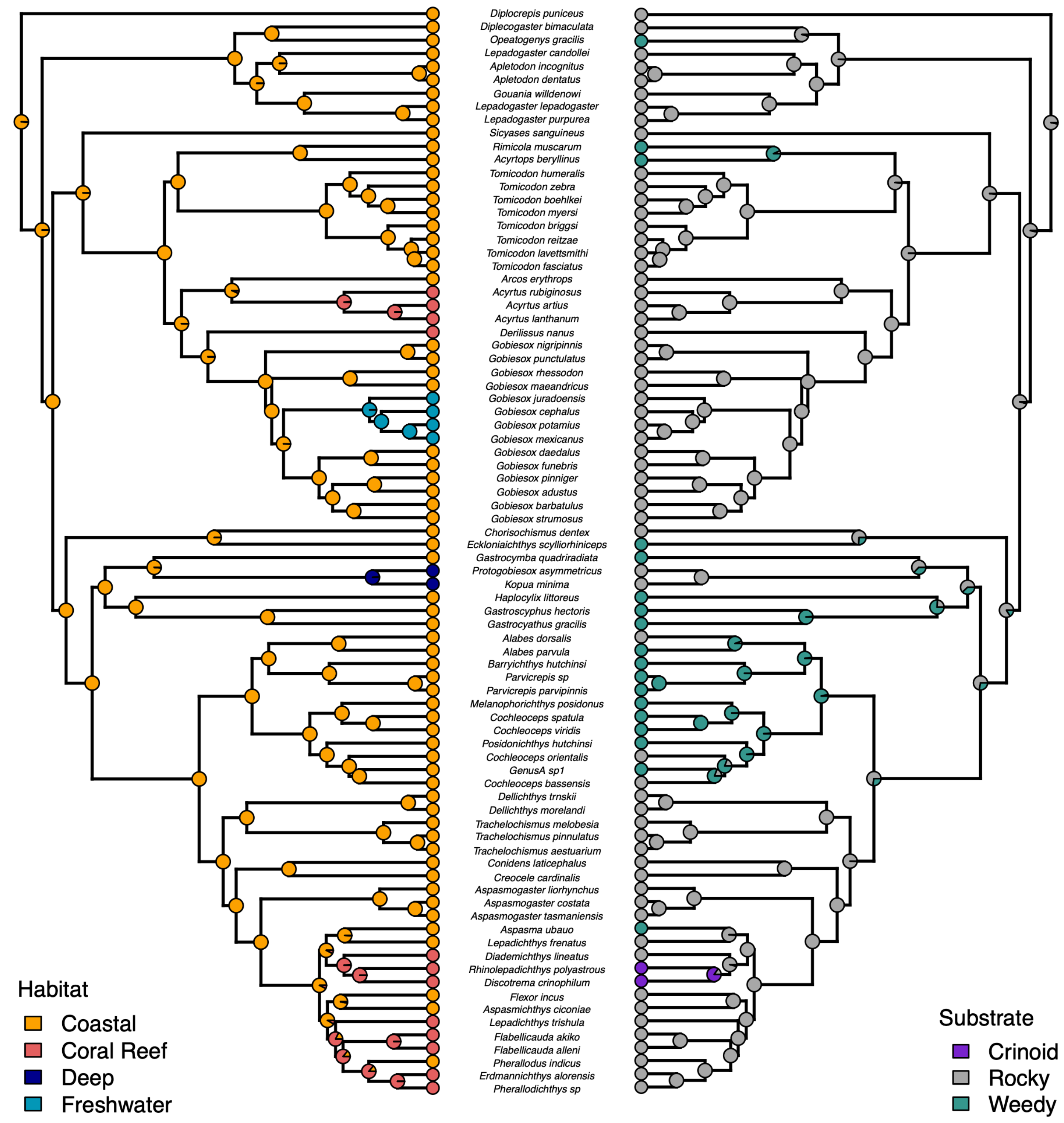


**Figure S2.** Ancestral state reconstructions of habitat and substrate use. The evolution of habitat and substrate was jointly reconstructed using corHMM, but the results for each trait are shown separately.
